# Antigen-specific competitive inhibition of CD4^+^ T cell recruitment into the primary immune response

**DOI:** 10.1101/2020.08.31.276527

**Authors:** Alexandra J. Spencer, Adrian L. Smith, Barbara Fazekas de St Groth

**Affiliations:** Centenary Institute of Cancer Medicine and Cell Biology, Newtown NSW 2042, Australia; Faculty of Medicine and Health, University of Sydney, NSW 2006, Australia

**Keywords:** T Cells, Cell Activation, Cell Proliferation, Competition for Antigen

## Abstract

Previous studies suggest that recruitment of naïve T cells into a program of cell division and differentiation is a highly synchronous process under tight regulation. However it is not known whether antigen availability is the major regulator of this process, or whether other factors such as ongoing responses to unrelated antigens can affect the size of the primary response. We have developed an adoptive transfer system to investigate the efficiency with which additional antigen specific cells are recruited into an ongoing primary immune response. Recruitment of additional cells is an inverse function of the size of the response and is progressively inhibited with time. Cells recruited late into the response proliferate less, and fewer secrete IL-2 and IFN-γ. Thus the size of the response changes very little as a result of late recruitment. The inhibition of recruitment, proliferation and differentiation affects only cells of the same specificity as the ongoing response, indicating that the size of an antigen specific response is independent of any shared factors such as access to APCs, costimulation or cytokines. Thus, during infection, the immune system retains the ability to respond as necessary to secondary infections or antigens not presented until later stages of the response.

## Introduction

Recruitment of naïve T cells into immune responses occurs in secondary lymphoid organs where dendritic cells located in the T cell zones express the antigen-MHC complexes and costimulatory molecules required for full T cell activation and differentiation (1). When foreign antigen reaches a lymphoid organ, T cells of appropriate specificity may be recruited from the pool of cells already present in the organ, or from circulating T cells that enter the organ after the arrival of antigen. Recent in vivo imaging studies have visualized the latter process, using a model in which all responder T cells enter the lymph node more than 18 hours after antigen-bearing DCs (2, 3). However earlier studies of the antigen-specific response of CFSE labelled naïve T cells have indicated that T cell recruitment into cell division in draining lymph nodes is remarkably synchronous (4), suggesting that the period during which T cells can be recruited into cell division is restricted. In addition, responses to different T cell epitopes peak simultaneously in vivo (5-7) suggesting the existence of control mechanisms independent of TCR affinity and antigen availability. Mechanisms to prevent recruitment of specific T cells into an ongoing response are not well understood, nor is it clear whether such mechanisms function in an antigen-specific or non-specific manner, an issue with important implications for the ability of the immune system to mount responses to the complex mixtures of antigen released by infectious organisms and vaccine preparations.

To investigate this question, we studied the efficiency with which naïve antigen-specific CD4^+^ T cells reactive with moth cytochrome C (MCC) are recruited into an ongoing immune response, as a function of the delay between initiation of the response by a first cohort of MCC-specific TCR transgenic responder cells and adoptive transfer of a second cohort of cells of the same specificity. We found that recruitment into cell division and cytokine production decreased as the time delay between the two cell cohorts increased, leading to a significant drop in recruitment with a delay of only 24 hours. Injection of additional antigen could partially compensate for this decline, suggesting that lack of available antigen limited recruitment of specific cells trafficking to the node after the initiation of the response. A role for antigen non-specific factors such as access to APCs, costimulatory signals or cytokines was ruled out by showing that the response to a second, independent antigen was unaffected by an ongoing response, even when the same DCs were presenting both antigens. These data illustrate the mechanism whereby the immune system responds efficiently to late phase antigens or secondary infections, while exerting tight control over the size and kinetics of each individual antigen specific response.

## Methods

### Experimental Animals

All lines of transgenic mice were bred and housed under specific pathogen-free conditions in the Centenary Institute Animal Facility. Approval for all animal experimentation was obtained from the Institutional Ethics Committee at the University of Sydney. 5C.C7 TCR transgenic (8, 9) and 3A9 TCR transgenic (10) mice were maintained on a B10.BR (H-2^k^) background and were crossed with either C57BL/6 or B6.SJL^Ptprca^ mice (Animal Resources Centre, Perth, Australia), to generate either CD45.2/CD45.2 homozygous or CD45.1/CD45.2 F1 progeny for use in experiments.

### Antigens and Immunisation

MCC peptide 87-103 (KANERADLIAYLKQATK) and HEL peptide 46-61 (NTDGSTDYGILQINSR) were purchased from Auspep Pty. Ltd. (Parkville, Vic, Australia). Mice were immunised subcutaneously with a mixture of peptides in PBS, emulsified in Complete Freunds Adjuvant (CFA) (Sigma Aldrich, St Louis, MI). Mice received a total volume of 200μl peptide-CFA emulsion distributed between injection sites in both hind footpads (50μl each) and the base of the tail (100μl). In mice that received two separate injections of antigen in CFA, 100ul was distributed between the two-hind pads for the first injection and a second 100ul injection at the base of the tail 4 days later.

### Expression and purification of recombinant HELMCC protein

MCC protein was synthesized in recombinant form because it is not available commercially. Initially the C-terminal MCC epitope was fused to the C-terminal end of hen egg lysozyme (HEL). However published studies have indicated that when the C-terminal MCC epitope was fused to the C-terminal end of GST, the MCC epitope could be presented to transgenic T cells without a requirement for processing, suggesting lack of secondary structure (11). Therefore to ensure that the presentation of both HEL and MCC antigens required processing, a modification of the system of Verma (12) was adopted. Molecular modelling of a number of lysozyme proteins derived from different bird species was used to identify a hypervariable loop flanked by conserved cysteine residues that was unlikely to be required for protein folding and stability. In the HEL protein, this loop corresponded to the 11 amino acids between cysteine residues 64 and 76 of the mature protein (residues 82 and 94 when the numbering begins at the start of the signal sequence). To make the HELMCC fusion protein, these 11 aa were replaced by a total of 19 aa encoding the C-terminal sequence of MCC (residues 87-103, KANERADLIAYLKQATK) flanked by an additional lysine at either end. This created two cathepsin cleavage sites on either side of the MCC sequence, previously shown to increase the efficiency of MCC processing and presentation (12). The original HELMCC sequence was generated by PCR amplification of a HEL cDNA (13) in two sections, each containing half of the MCC sequence linked by a naturally occurring *Bgl II* site at position 21 of the 57bp sequence. The primers were: for the left arm, 5’ primer TAGGGATCCATGAGGTCTTTGCTAATCTTGGTG, 3’primer TAGCAGATCTGCGCGTTCGTTGCGTTTTTTGCACCACCAGCGGCT; for the right arm, 5’ primer GCAGATCTGATTGCGTATTTAAAACAGGCAACCAAAAAATGCAACATCCCGTGC, 3’primer TAGATGGAATTGTCTGCAGCCCAGCCGGCAGCCTCT. The final contruct was cloned into pHFBgl vector (Armen B. Shanafelt, DNAX research Institiute San Alto, CA, USA) between the BamHI and Pst I sites for expression of recombinant HELMCC protein in insect cells. HELMCC was subsequently cloned into pcDNA3 (Invitogen) for transfection of CHO cells to test protein expression and antigenicity. This expression system yielded minimal amouts of protein and therefore a yeast expression system was adopted, with the assistance of Didrik Paus and Robert Brink. To enable expression in the yeast system, the Bgl II site was first removed by site-directed mutagenesis. A C-terminal His(6)-tag (SGHHHHHH) was added using the primers 5’ GACCGTCTCGAGAAAAGAAAAGTCTTTGGACGATGTGAGC and 3’ GTGACTGAATTCTTACTAATGGTGATGGTGGTGATGGCCGCTCAGCCGGCAGCCTCTGA TCCAC and the product was subsequently cloned into the vector pPIC9K (Invitrogen) between the Xho I and EcoR I sites by doing a three-way ligation (EcoR I, Xho I and Pci I) as the Xho I site is not unique. The vector was linearized with BglII and transformation of yeast (Pichia pastoris) and induction of recombinant protein expression was carried out according to the manufacturer’s instructions (Invitrogen). Recombinant HELMCC protein was purified from yeast culture supernatants by ionexchange chromatography using Ni^2+^ affinity chromatography (Amersham Biosciences). Recombinant HELMCC protein was >90% pure as determined by SDS-PAGE and staining with Coomassie blue. The dose of HELMCC was chosen after performing in vivo dose response experiments, using comparison to MCC and HEL peptide dose response curves.

### Adoptive Transfer

Pooled unfractionated inguinal, axillary, subscapular and para-aortic lymph nodes served as the source of donor TCR transgenic T cells for adoptive transfer. Transgenic lymph node cells were labelled with 5-carboxyflurorescein diacetate succinimidyl ester (CFSE) (Molecular Probes) as described (14) prior to transfer into recipient mice. The percentage of CD4^+^ cells in the lymph node cell preparation was determined by flow cytometry before injection and the total number of cells adjusted to ensure each recipient mouse received the same number of CD4^+^ TCR transgenic T cells at each timepoints. Unless otherwise noted, each cell injection contained 2.5×10^6^ CD4^+^ TCR transgenic T cells.

### Flow Cytometry

To distinguish CFSE-labelled CD45.2 homozygous donor cells from host and competing cells expressing CD45.1, anti-CD45.1 mAb (A20.1) was conjugated in-house with either allophycocyanin (Molecular Probes) or Pacific Blue (Invitrogen). CFSE labelled cells were detected with either anti-CD4-allophycocyanin (EBiosciences, San Diego, CA) or anti-CD4-PE (BD, Franklin Lakes NJ). Anti-CD69-PE (BD) was used to measure CD69 expression. Intracellular cytokines were detected with either anti-IL-2-PE (BD) or anti-IFN-γ-PE (BD) and samples compared to control samples stained with isotype control mAbs IgG2a-PE or IgG1-PE. Dendritic cells were stained with a combination of CD11c-PE (BD), B220-PerCP (BD) and anti-IE^k^/IA^k^ (M5-114) conjugated to allophycocyanin (Ebiosciences). Dead cells were excluded using either 2-(4-Aminophenyl)-6-indolecarbamidine dihydrochloride (Sigma Aldrich) or Propidium Iodide (Sigma Aldrich). In some experiments, transgenic cells were also identified with a biotinylated mAb against one of the two TCR chains. 5C.C7 cells were identified with a mAb against either Vα11 (RR8.1) or Vβ3 (KJ25) and 3A9 cells with a mAb against Vβ8.2 (F23.2). 5 or 6 colour flow cytometric analysis was performed using either a FacsCalibur, LSRII or FacsVantage machine (Beckton Dickenson, San Jose CA).

### Antigen restimulation and intracellular cytokine staining

1×10^7^ draining lymph node or spleen cells were restimulated for 6 hours with 10μM MCC peptide with the addition of 5μM Brefeldin A (Sigma Aldrich) for the final 4 hours. Following stimulation, cells were stained for surface molecules prior to fixation in 4% paraformadehyde (Sigma Aldrich) for 20 minutes. Cells were permeablised by washing in saponin/BSA solution (0.5% saponin and 0.1% BSA) prior to staining for intracellular cytokines with mAbs diluted in saponin/BSA solution.

### Analysis of flow cytometric data

Data were analysed using FlowJo Software (TreeStar, Ashland, OR). The CFSE profile of dividing cells was analysed as described earlier (4, 15). Briefly, to calculate the percentage of cells recruited into cell division, the number of cells in each cell division peak was corrected for the increase due to division itself. Thus, the number of cells in the first division peak was divided by 2, the second by 4 and so on. The percentage of cells recruited into cell division (R) was calculated using Equation 1: 

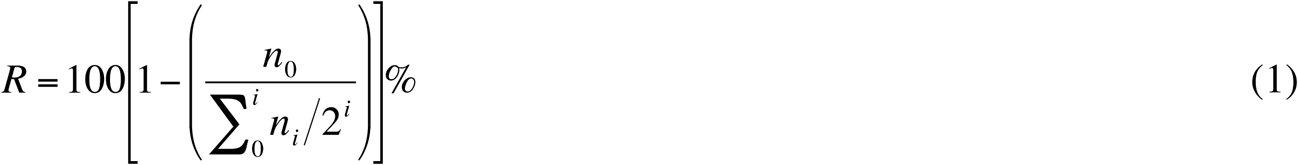

where *n*_*i*_ = the number of cells in the *i*th division peak.

The percentage of divided cells in the *i*th division peak (D_*i*_) was calculated using Equation 2: 

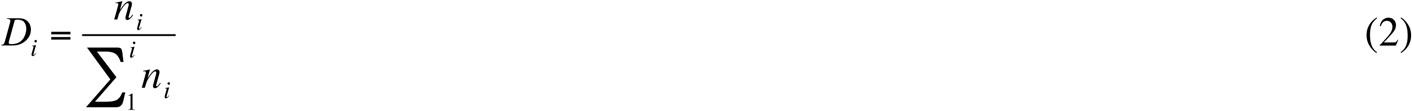

## Results

### An ongoing response progressively inhibits the recruitment of additional precursors

Recruitment of naïve T cells into an ongoing response was measured in the experiment described in Figure 1. The treatment of the 3 experimental groups is detailed in Figure 1a. All recipient mice were immunised subcutaneously with 10μg MCC peptide in CFA and received an adoptively transferred cohort of CFSE-labelled 5C.C7 cells 1.5 days later. In the control group (i), no other 5C.C7 cells were administered. Group (ii) received an equal number of unlabelled 5C.C7 cells at the same time as the CFSE-labelled cohort, whereas in group (iii), the 5C.C7 response was initiated by transfer of an unlabelled cohort of 5C.C7 cells 1 day before the CFSE-labelled cells. The response of the CFSE-labelled cohort was analysed 2.5 days after adoptive transfer, when cells proliferating at a maximal rate were still distinguishable from unlabeled cells.

**Figure 1:**
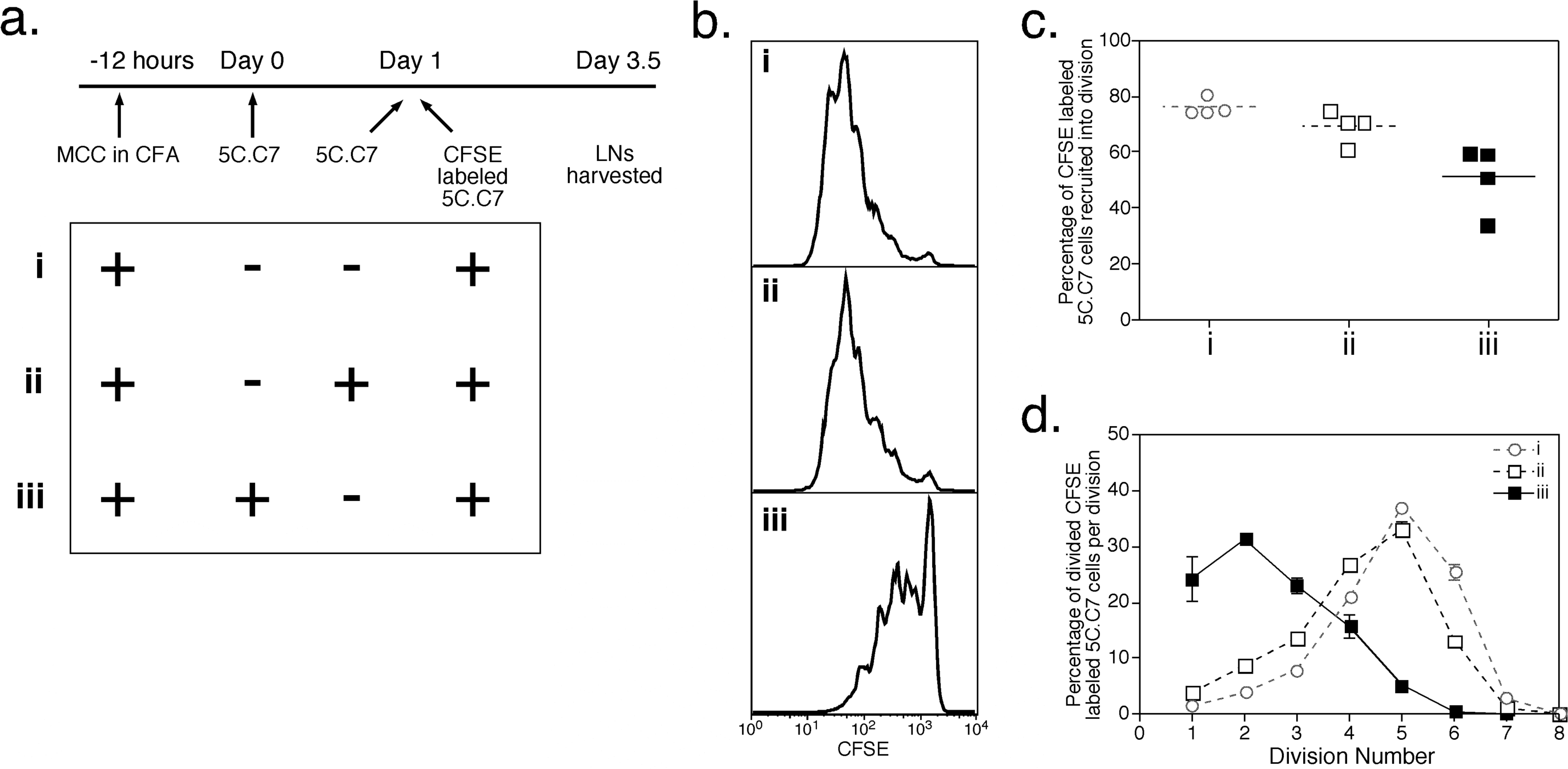
An in vivo model to study recruitment of CD4^+^ T cells into an ongoing response. (a) Experimental plan. Three groups of 4 recipient mice were immunized with 10μg MCC peptide in CFA and 1.5 days later were adoptively transferred with CFSE-labelled 5C.C7 TCR transgenic lymph node cells containing 2.5×10^6^ CD4^+^ T cells. In the control group (i), no other cells were administered. Group (ii) received an equal number of 5C.C7 cells at the same time as the CFSE-labelled cohort, whereas group (iii) received an equal number of 5C.C7 cells 1 day before the CFSE-labelled cohort. Draining lymph nodes were harvested 2.5 days after the CFSE-labelled adoptive transfer. (b) Representative CFSE profiles of donor 5C.C7 cells, gated for live CD4^+^ T cells expressing the transgenic TCR Vβ11 (KJ25). (c) Percentage of CFSE-labelled CD4^+^ TCR transgenic T cells recruited into cell division, calculated as described in Materials and Methods. Each point represents an individual mouse and the group mean is indicated by the horizontal line. (d) Percentage of divided CFSE-labelled CD4^+^ TCR transgenic T cells in each division peak, calculated as described in Materials and Methods. The number of cells in each division peak was divided by the total number of divided cells (ie, excluding undivided cells). Each point represents the mean calculated from the 4 mice in each group and the error bars show the SEM.

In the absence of a competing cohort of 5C.C7 cells, 80% of CFSE-labelled 5C.C7 cells in the draining lymph nodes were recruited into cell division and the majority of divided cells progressed through 4-6 divisions in 2.5 days (Fig. 1b-d, group (i)). Under these circumstances, a prior response of polyclonal host T cells would also be initiated by MCC-CFA immunisation, but clearly was not sufficiently strong to prevent a robust response by the cohort of high affinity 5C.C7 cells.

Simultaneous transfer of an equal number of competing 5C.C7 cells (group (ii)) led to a slight but not statistically significant reduction in recruitment without a significant change in the CFSE division profile (Fig. 1b-d, compare groups (i) and (ii)). This result is consistent with our previously published data showing that a 2-fold increase in high affinity precursor frequency at the initiation of the primary response does not significantly reduce recruitment on a per cell basis (16). However when a competing high affinity response was initiated by transfer of 5C.C7 cells 1 day earlier, recruitment into division was reduced from 76% to 51% and the majority of divided cells passed through only 1-3 divisions in 2.5 days (Fig 1b-d, compare groups (i) and (iii)). Thus a delay of only 24 hours had a major impact on the ability of T cells to participate in an ongoing high affinity response to antigen.

In a second experiment, outlined in Fig 2a, a longer delay was introduced between the initiation of the competing high affinity response and the transfer of CFSE-labelled cells. To do this, the CFSE-labelled cell transfer was delayed until 3.5 days after immunisation. This did not significantly reduce the control response (compare groups (i) in Figs 1 and 2), indicating that the degree of competition exerted by an ongoing response of host polyclonal T cells was far smaller than that of a cohort of high affinity 5C.C7 cells. Increasing the delay between initiation of the 5C.C7 response and administration of the CFSE-labelled cohort of 5C.C7 cells from 1 day to 3 days (groups (ii) and (iii), Fig.2b-d) reduced recruitment and cell division markedly, with fewer than 5% of cells recruited into the response and, of these, 50% progressing through only a single round of cell division in 2.5 days.

**Figure 2:**
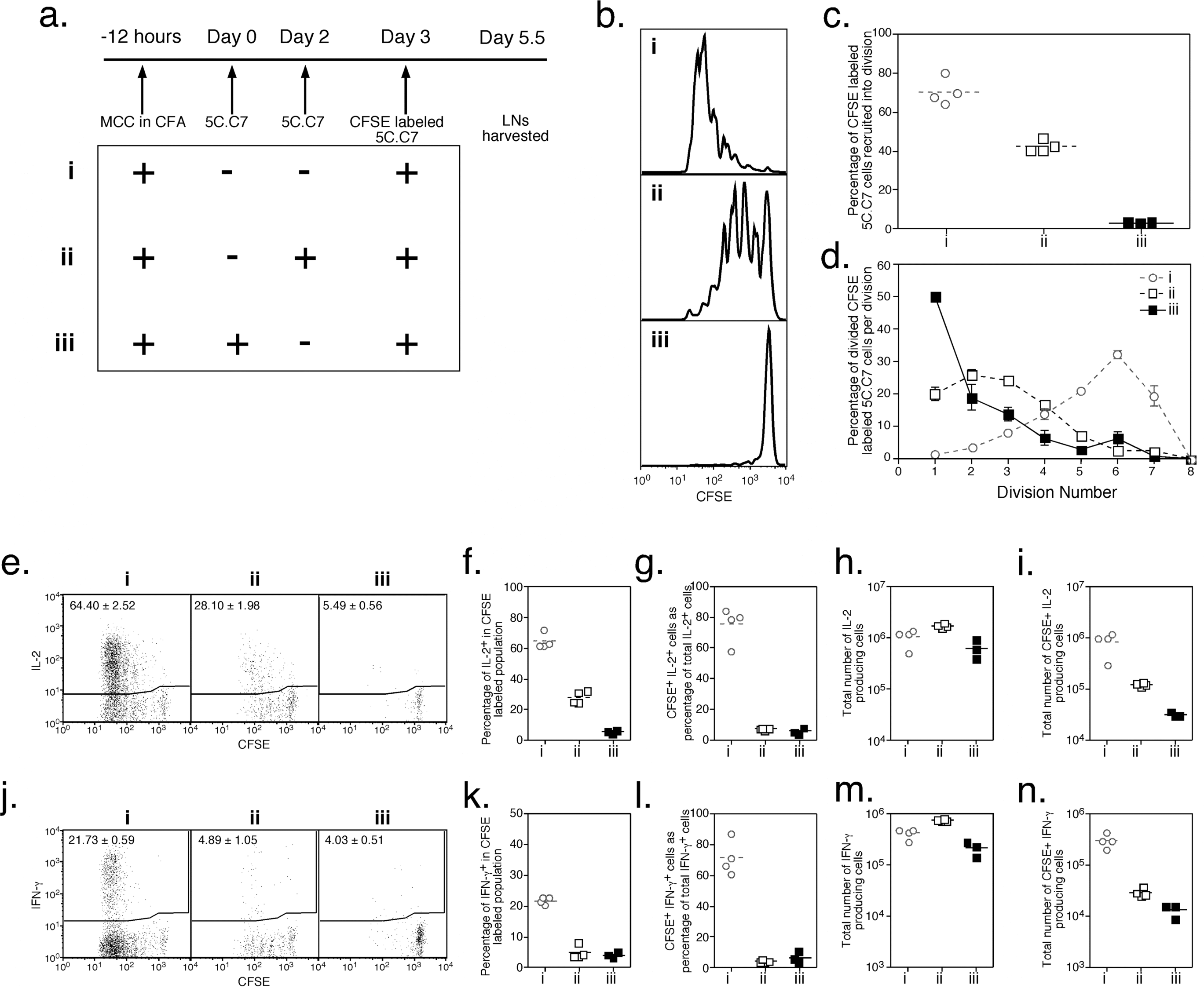
Ongoing responses progressively inhibit the recruitment of naive CD4^+^ cells. (a) Experimental plan. Three groups of 4 recipient mice were immunized with 10μg MCC peptide in CFA and 3.5 days later were adoptively transferred with CFSE-labelled 5C.C7 TCR transgenic lymph node cells containing 2.5×10^6^ CD4^+^ T cells. In the control group (i), no other cells were administered. Groups (ii) and (iii) received an equal number of 5C.C7 cells 1 or 3 days before the CFSE-labelled cohort. Draining lymph nodes were harvested 2.5 days after the CFSE-labelled adoptive transfer. (b) Representative CFSE profiles of donor 5C.C7 cells, gated for live CD4^+^ T cells expressing the transgenic TCR Vα11 (RR8). (c) Percentage of CFSE-labelled TCR transgenic CD4^+^ T cells recruited into cell division. Each point represents an individual mouse and the group mean is indicated by the horizontal line. (d) Percentage of divided CFSE-labelled TCR transgenic CD4^+^ T cells in each division peak. Each point represents the mean calculated from the 4 mice in each group and the error bars show the SEM. (e-n) Production of IL-2 and IFN-γ, measured by intracellular cytokine staining after 6 hours of ex vivo restimulation with 10μM MCC peptide and Brefeldin A for the final 4 hours. (e, j) Representative dot plots of CFSE vs intracellular IL-2 or IFN-γ in CFSE-labelled TCR transgenic 5C.C7 cells, gated for live CD4^+^ T cells. The gates were calculated on the basis of isotype control staining, as described in Materials and Methods. Numbers within dot plots represent the mean percentage +/-SEM of cells within the positive gate for each group. (f, k) Percentage of cytokine-producing cells, calculated from the profiles as in (e, j). (g, l) Percentage of total cytokine-producing cells derived from the CFSE-labelled cohort of CD4^+^ TCR transgenic T cells, calculated by dividing the number of CD4^+^ TCR transgenic CFSE-labelled cytokine-producing cells in each sample by the total number of CD4^+^ cytokine-producing cells in the sample. (h, m) Total CD4^+^ cytokine-producing cells in the draining lymph nodes of each mouse, calculated by multiplying the percentage of CD4^+^ cytokine-producing cells in each sample by the number of cells harvested from the draining lymph nodes. (i, n) Total CFSE-labelled TCR transgenic CD4^+^ cytokine-producing cells in the draining lymph nodes of each mouse, calculated by multiplying the percentage of CFSE-labelled TCR transgenic CD4^+^ cytokine-producing cells in each sample by the number of cells harvested from the draining lymph nodes. For panels (e-i) and (k-n), each point represents an individual mouse and the group mean is indicated by the horizontal line.

In the same experiment, the ability of late-arriving precursors to contribute to the overall effector response was measured by intracellular cytokine staining after 6 hours of ex vivo restimulation. For cells competing against a cohort injected 1 day before, the frequency of IL-2-producing cells was reduced to 28% compared with 64% for the control group (compare groups (i) and (ii), Fig. 2e-f). IFN-γ-producing 5C.C7 cells were also reduced from 22% to 5% (Fig. 2j-k). An even smaller percentage of cells competing against a cohort injected 3 days earlier was positive for intracellular IL-2 or IFN-γ. To estimate the contribution of the CFSE-labelled cohort of cells to the total IL-2 and IFN-γ responses, the number of cytokine-producing cells derived from the CFSE-labelled cohort was calculated as a percentage of total IL-2-or IFN-γ-producing cells (Figs 2g, l). In groups and (iii), the contribution of these late arriving cells to the total IL-2 and IFN-γ responses was less than 5%, whether competing cells were transferred 1 or 3 days previously (Fig. 2g, k). In contrast, the CFSE-labelled high affinity cohort transferred 3.5 days after immunisation contributed 60-80% of IL-2- and IFN-γ-producing cells, the remaining 20-40% being derived from the polyclonal host repertoire (Fig. 2g, k, group (i)). This result indicates that endogenous cells could contribute significantly to the effector cell response when injection of a competing cohort was delayed, but even then they did not contribute the majority of effector cells. The size of the total MCC specific cytokine response (Fig. 2h, m) was similar for all 3 groups, the slight decrease in the case of group (iii) being attributable to the kinetics of cytokine production in this model. We have previously shown that the number of cells secreting IFN-γ and IL-2 in the draining node in response to peptide-CFA peaks at day 4 after immunisation (Bugeja and Fazekas de St. Groth, unpublished data). Consistent with this, the highest absolute number of IL-2 or IFN-γ secreting cells was present after restimulation of cells from group (ii), in which the majority of responder cells were injected 3.5 days before harvest, rather than in group (i) (2.5 days before harvest) or group (iii) (5.5 days before harvest).

Taken together, these data indicate that recruitment of naïve antigen-specific precursors decreased over the course of a primary immune response. Their contribution to the effector cell pool was inversely related to the size of the ongoing response. Thus 2.5×10^6^ 5C.C7 cells generated 10^6^ IL-2-producing and 3×10^5^ IFN-γ-producing cells when a polyclonal host response was initiated 3 days earlier (group (i), Fig. 2i, n), but only 3 ×10^4^ IL-2-producing and 10^4^ IFN-γ-producing cells when the response was initiated 3 days earlier by 2.5×10^6^ 5C.C7 cells (group (iii), Fig. 2i, n).

### Additional antigen can enhance recruitment into an ongoing response

To test whether competition for access to specific antigen was responsible for exclusion of antigen-specific T cells from an ongoing response, additional antigen was administered at the time of CFSE-labelled responder cell transfer, according to the schema presented in Fig 3a. In this experiment, a 4 day delay was introduced between the initiation of the response and the transfer of CFSE-labelled cells. Consistent with the results presented above, both recruitment and ongoing cell division were reduced by initiation of a high affinity response 4 days earlier (compare groups (i) and (iii), Fig. 3b-d). However administration of additional antigen at the time of CFSE-labelled cell transfer restored recruitment to over 80% (group (iv), Fig. 3b,c), although cell division was only partially restored (Fig. 3d). These data suggested that restricted access to antigen was at least partially responsible for the failure to recruit newly arrived cells during an ongoing response.

**Figure 3:**
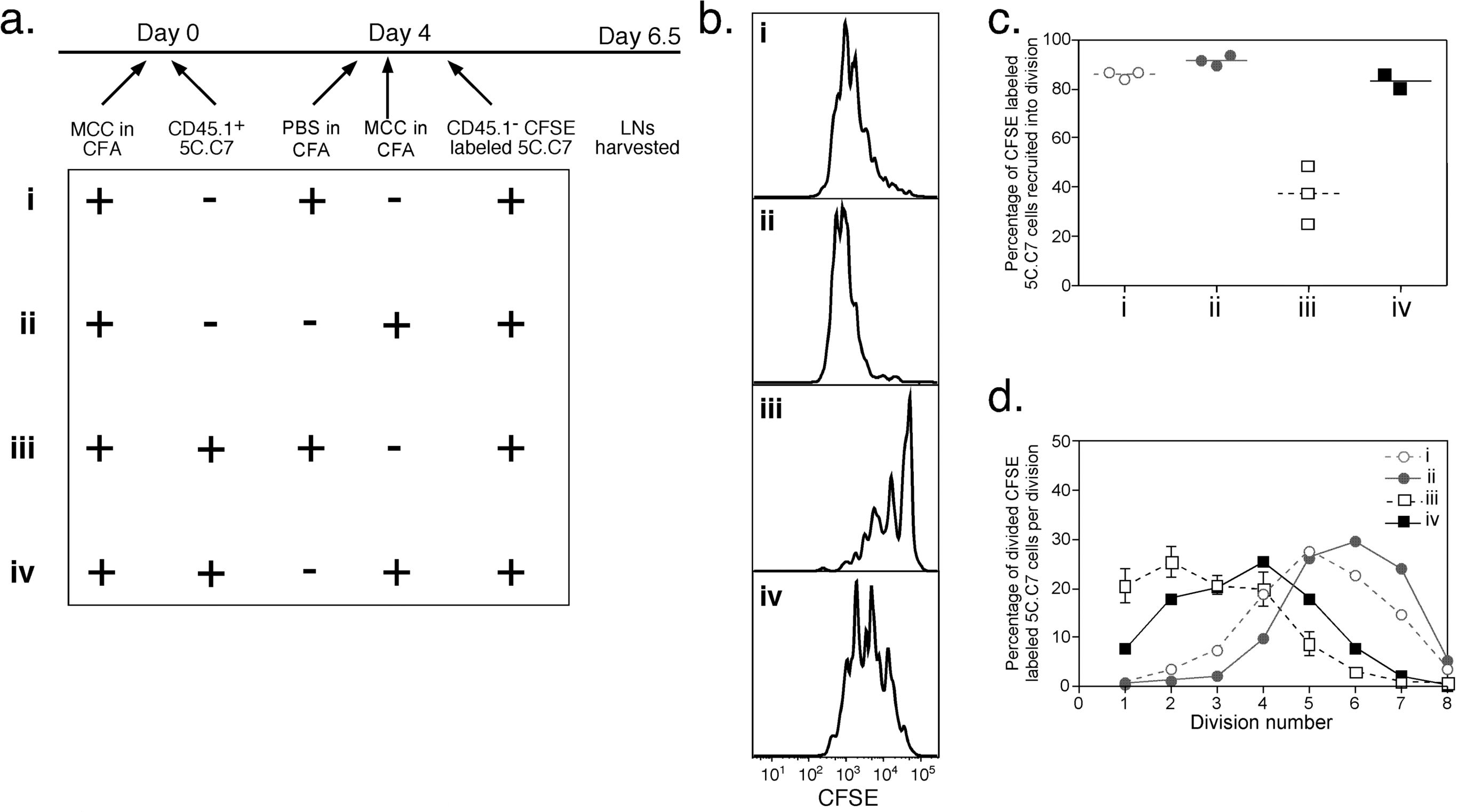
Addition of more antigen enhances recruitment of naive CD4^+^ T cells into an ongoing response. (a) Experimental plan. Four groups of 4 CD45.1^+^ recipient mice were immunized with 10μg MCC peptide in CFA and 4 days later were adoptively transferred with CFSE-labelled CD45.1^-^ 5C.C7 TCR transgenic lymph node cells containing 2.5×10^6^ CD4^+^ T cells. The control group (i) was immunised with PBS/CFA at the time of adoptive transfer, whereas group (ii) received a further injection of MCC/CFA. Groups (iii) and (iv) received an equal number of CD45.1^+^ 5C.C7 cells 4 days before the CFSE-labelled cohort, and were immunised with PBS/CFA or MCC/CFA, respectively, at the time of the CFSE-labelled adoptive transfer. Draining lymph nodes were harvested 2.5 days later. (b) Representative CFSE profiles of donor CD45.1^-^ 5C.C7 cells, gated for live CD4^+^ T cells expressing Vα11. (c) Percentage of CFSE-labelled TCR transgenic CD4^+^ T cells recruited into cell division. Each point represents an individual mouse and the group mean is indicated by the horizontal line. (d) Percentage of divided CFSE-labelled TCR transgenic CD4^+^ T cells in each division peak. Each point represents the mean calculated from the 4 mice in each group and the error bars show the SEM.

### Continuous competition for peptide-MHC limits the size of the CD4 response

As a direct measure of access to antigen by newly transferred 5C.C7 T cells, CD69 expression was determined 2 hrs after adoptive transfer (Fig. 4). In this experiment, the first and second cohorts of TCR transgenic cells were distinguished by expression of different CD45 alleles. The experimental groups are detailed in Fig. 4a, and comprised mice that had been injected with 5C.C7 T cells 1, 2 or 3 days before the CFSE-labeled cohort, in addition to the control group (i) in which no 5C.C7 cells were injected previously. Group (v) was included to control for the antigen-specificity of the ongoing primary immune response. In this group, the initial response of adoptively transferred 3A9 TCR transgenic cells was directed to a non-cross-reactive antigen, HEL, presented on a different MHC molecule (IA^k^). The choice of peptide-MHC specificity was designed to exclude any competition between peptides for presentation by APCs. All recipients were immunised with a mixture of MCC and HEL peptides, the doses chosen to give equivalent cell division profiles of 5C.C7 and 3A9 TCR transgenic cells at day 2.5.

**Figure 4:**
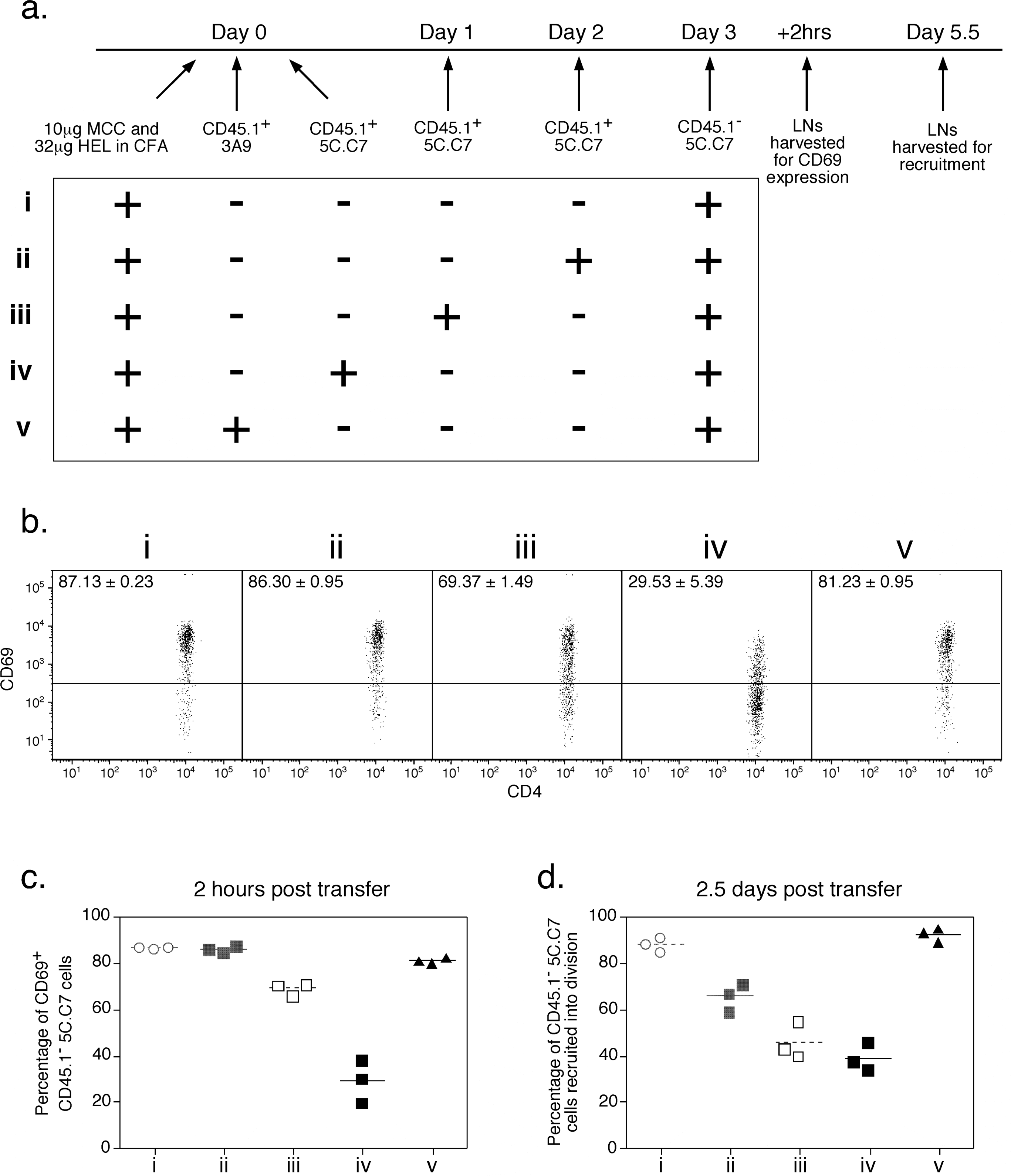
Recruitment into division as a function of early access to antigen presenting cells. (a) Experimental plan. Five groups of 6 CD45.1^+^ recipient mice were immunized with an emulsion containing 10μg MCC and 32μg HEL peptide in CFA. 3 days later, all mice were adoptively transferred with CFSE-labelled CD45.1^-^ 5C.C7 TCR transgenic lymph node cells containing 2.5×10^6^ CD4^+^ T cells. Groups (ii) to (iv) received an equal number of CD45.1^+^ 5C.C7 cells 1, 2 or 3 days before the CFSE-labelled cohort. Group (v) received CD45.1^+^ 3A9 TCR transgenic lymph node cells containing 2.5×10^6^ CD4^+^ T cells 3 days before the CFSE-labelled cohort. Draining lymph nodes from 3 mice per group were harvested 2 hours after adoptive transfer of the CFSE-labelled cohort to measure CD69 upregulation, and the remaining 3 mice per group were sacrificed 2.5 days later to calculate recruitment into cell division. (b) Representative dot plots of CD4 vs CD69 in CD45.1^-^ CD4^+^ 5C.C7 TCR transgenic draining lymph node cells 2 hours after adoptive transfer. Numbers within dot plots represent the mean percentage +/-SEM of cells within the positive gate for each group. (c) Percentage of CFSE-labelled 5C.C7 TCR expressing CD69 calculated from the plots in (b). Each point represents an individual mouse and the group mean is indicated by the horizontal line. (d) Percentage of CFSE-labelled TCR transgenic CD4^+^ T cells recruited into cell division. Each point represents an individual mouse and the group mean is indicated by the horizontal line.

In the control group (i), the percentage of cells that upregulated CD69 expression within the first 2 hours after transfer corresponded to the percentage recruited into division (Fig. 4b). Thus 87% of antigen-specific T cells upregulated CD69 and 88% entered cell division. However as the delay between the initiation of the high affinity response and the transfer of the CFSE-labelled cohort increased, it became apparent that not every cell expressing CD69 was recruited into division. With a 1 day delay (group (ii)), over 80% of cells upregulated CD69 but a mean of only 69% were recruited into division. The disparity was even greater when 5C.C7 cells were transferred 2 days into an ongoing response. When the delay was 3 days, both CD69 expression and recruitment were reduced to less than 40%.

In contrast to the effect of a 4 day delay between adoptive transfer of cohorts of 5C.C7 cells, initial transfer of a cohort of 3A9 anti-HEL cells had no effect on the subsequent response of 5C.C7 cells (Fig. 4b-d, group (v)). Despite a minor but not statistically significant drop in CD69 upregulation (from 87% to 81%) when 5C.C7 cells were transferred 3 days into an ongoing 3A9 response, a similar percentage of 5C.C7 cells were recruited into division by day 5.5. This suggests that reduced recruitment of naïve cells into an ongoing response is due to antigen-specific factors rather than a general inability to access APCs or co-stimulatory signals, as suggested previously (17, 18).

### Activation of CD4 T cells is unaffected by ongoing responses to independent antigens

It is possible that an antigen non-specific competitive effect was masked in the experiments described above by the use of maximal doses of antigen. To test this, we reduced the peptide doses so that only half the antigen-specific T cells were recruited in the control group, and measured the effect of an ongoing 5C.C7 response on the lower affinity 3A9 response to HEL, rather than vice versa (Fig. 5). Even under these conditions, an ongoing 5C.C7 response was not capable of reducing recruitment and proliferation of a cohort of 3A9 cells adoptively transferred 3 days later and analysed after 2.5 days in vivo (Fig. 5b-d). In contrast, an ongoing 3A9 response profoundly decreased the response of a CFSE-labeled cohort of 3A9 cells adoptively transferred 3 days later. To test whether an effect of antigen non-specific competition was apparent later in the response, the number of 3A9 cells recovered from the draining nodes and spleens at day 10 of the response was determined in mice with and without an ongoing response of 5C.C7 cells initiated 1 day before transfer of the 3A9 cells (Fig. 6). Once again no effect of a response to an independent antigen could be detected (Fig. 6b-c).

**Figure 5:**
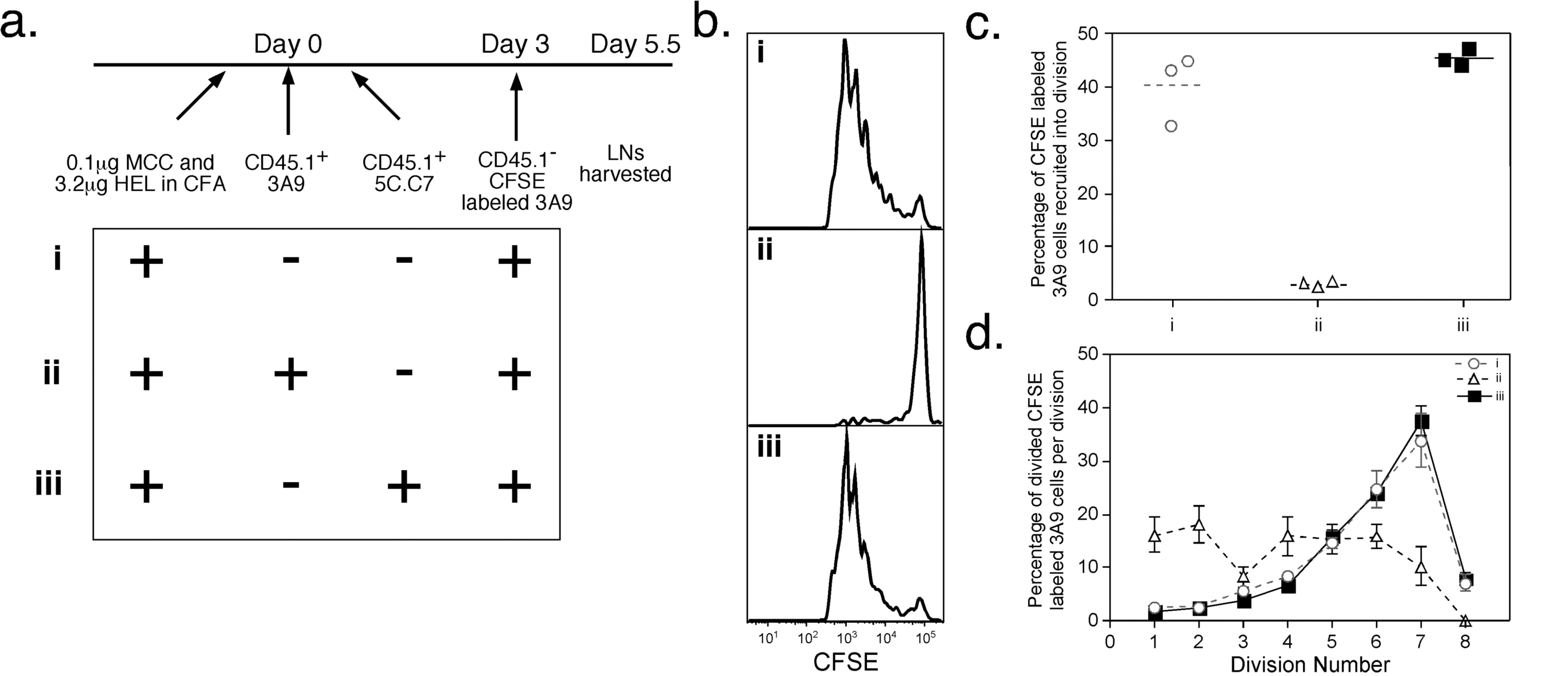
Lack of competition between simultaneous responses to unrelated antigens. (a) Experimental plan. Three groups of 4 CD45.1^+^ recipient mice were immunized with a mixture of 0.1μg MCC peptide and 3.2μg HEL peptide in CFA, and 3 days later were adoptively transferred with CFSE-labelled CD45.1^-^ 3A9 TCR transgenic lymph node cells containing 1×10^6^ CD4^+^ T cells. Groups (ii) and (iii) received an equal number of CD45.1^-^ 3A9 or 5C.C7 TCR transgenic cells 3 days before the CFSE-labelled cohort. Draining lymph nodes were harvested 2.5 days after the CFSE-labelled adoptive transfer. (b) Representative CFSE profiles of donor 3A9 cells, gated for live CFSE^+^ CD4^+^ T cells. (c) Percentage of CFSE-labelled 3A9 TCR transgenic CD4^+^ T cells recruited into cell division. Each point represents an individual mouse and the group mean is indicated by the horizontal line. (d) Percentage of divided CFSE-labelled 3A9 TCR transgenic CD4^+^ T cells in each division peak. Each point represents the mean calculated from the 4 mice in each group and the error bars show the SEM.

**Figure 6:**
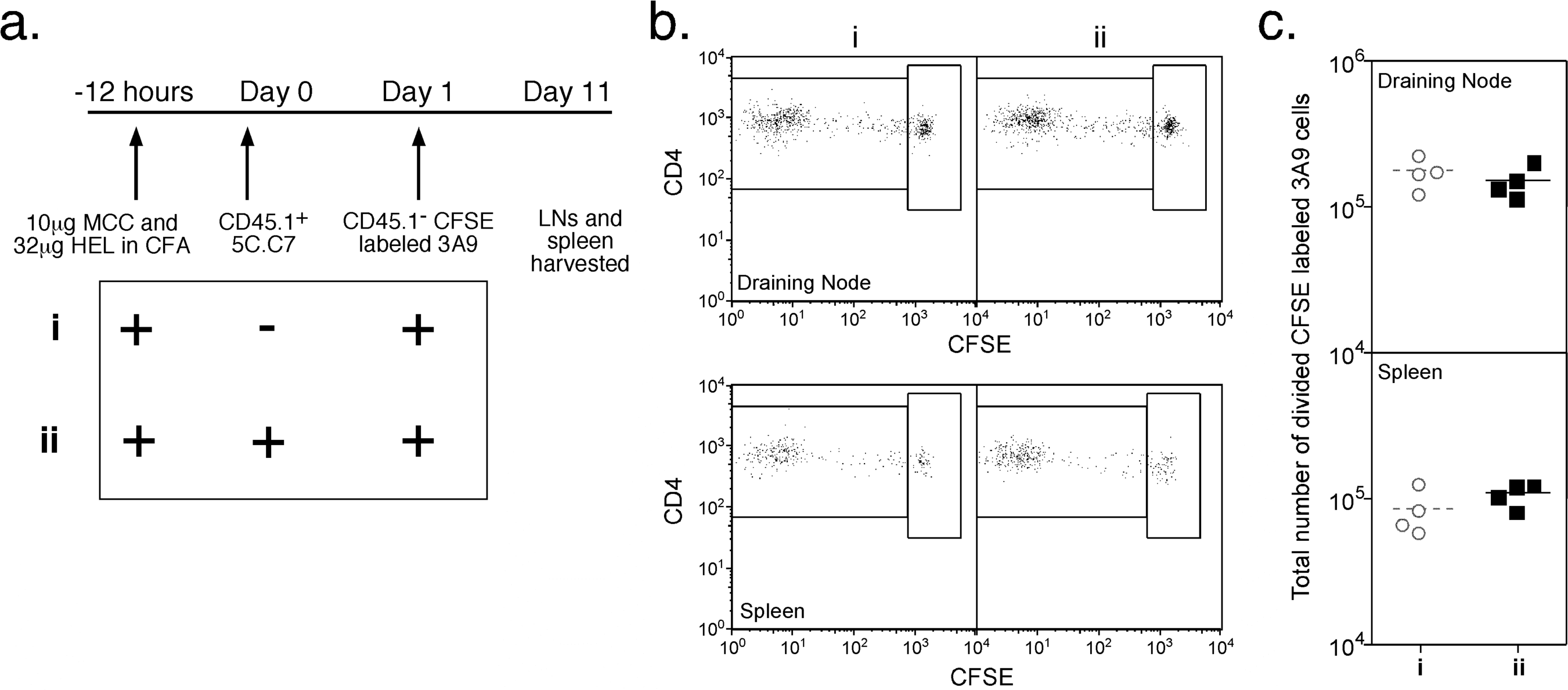
Day 10 responses in the presence of a simultaneous response to an unrelated antigeny. (a) Experimental plan. Two groups of 4 CD45.1^+^ recipient mice were immunized with a mixture of 10μg MCC peptide and 32μg HEL peptide in CFA. 1.5 days later, all mice were adoptively transferred with CFSE-labelled CD45.1^-^ 3A9 TCR transgenic lymph node cells containing 2.5×10^6^ CD4^+^ T cells. Group (ii) received an equal number of CD45.1^+^ 5C.C7 cells 1 day before the CFSE-labelled cohort. Draining lymph nodes and spleens were harvested 10 days after adoptive transfer of the CFSE-labelled cohort. (b) Representative dot plots of CFSE vs CD4 for CD45.1^-^ Vβ8.2^+^ CD4^+^ 3A9 TCR transgenic draining lymph node cells (upper panel) and spleen (lower panel). Gating for divided vs undivided 3A9 cells is shown. (c) Number of divided CD45.1^-^ CD4^+^ 3A9 TCR transgenic cells remaining in the draining lymph nodes after 10 days, calculated by multiplying the percentage of divided CD45.1^-^ CD4^+^ 3A9 TCR transgenic cells in each sample by the number of cells harvested from the draining lymph nodes. Each point represents an individual mouse and the group mean is indicated by the horizontal line.

These experiments strongly suggested that responses to non cross-reactive antigens were independent of each other. However, it remained theoretically possible that the responses to HEL and MCC were being driven by non-intersecting subsets of DCs, despite injection of the 2 peptides as a mixture emulsified in CFA. To overcome this problem, we stimulated transgenic cells with a recombinant protein, HELMCC, containing both the HEL and MCC epitopes. This protein requires processing for presentation of both epitopes, ensuring that both antigens would likely be presented at a fixed ratio by the same DCs (Fig. 7). The response of CFSE-labelled 3A9 cells (group (i)) was compared to that of 3A9 cells in mice with an ongoing 3A9 or 5C.C7 response (groups (ii) and (iii) respectively, Fig. 7a). Once again, only an ongoing response to the same antigen reduced recruitment and cell division of the labelled cohort (Fig. 7b-d). In a second experiment, the memory response of 3A9 cells transferred on day 3 of an ongoing 5C.C7 response was compared to a control 3A9 response by measuring the number of 3A9 cells 8 weeks after immunisation (Fig. 7e). Even at this late timepoint, there was no significant difference in the total number of 3A9 cells recovered from either the draining nodes or spleen in the 2 groups of mice. Thus we found no evidence of competition between cells of different antigen and MHC allele specificity at any point in the primary immune response.

**Figure 7:**
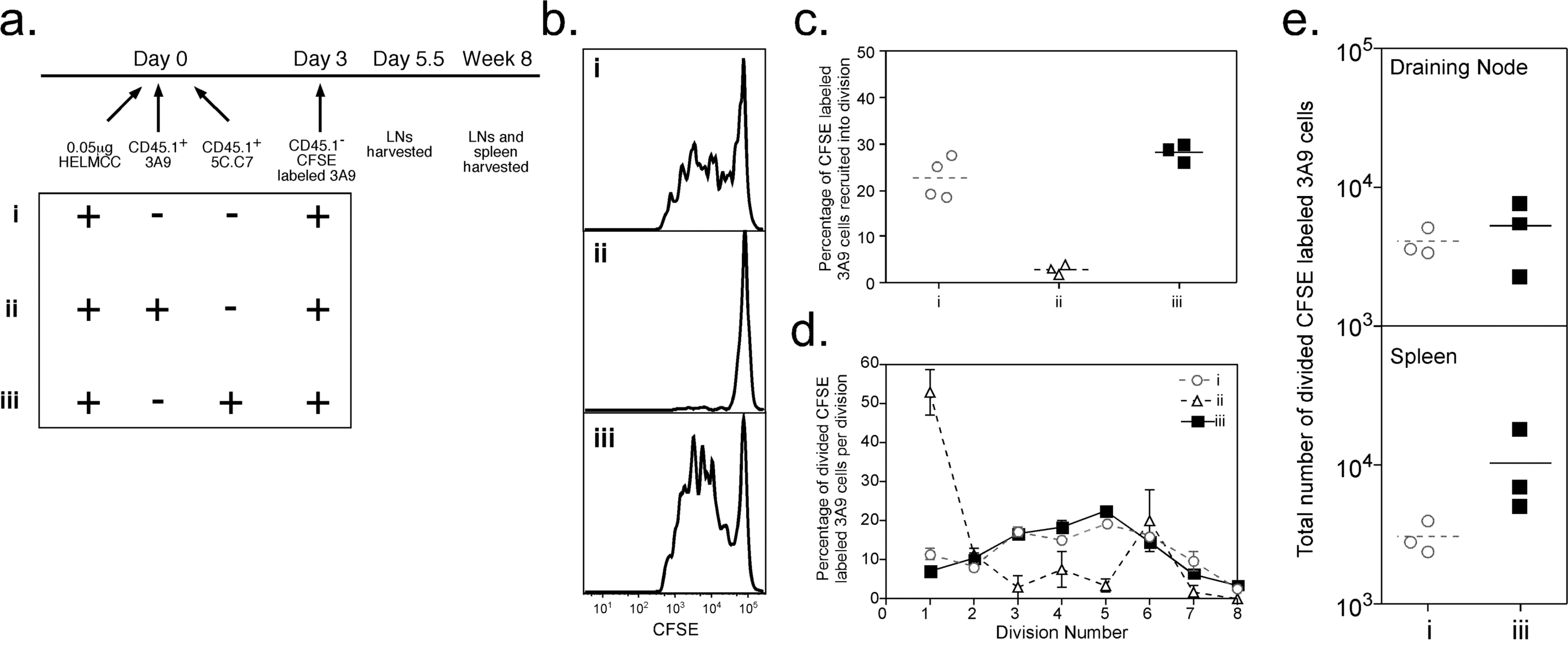
Presentation of two independent epitopes by the same APC. (a) Experimental plan. Three groups of CD45.1^+^ recipient mice were immunized with 0.5μg recombinant HELMCC in CFA. 3 days later, all mice were adoptively transferred with CFSE-labelled CD45.1^-^ 3A9 TCR transgenic lymph node cells containing 5×10^5^ CD4^+^ T cells. Groups (ii) and (iii) received an equal number of CD45.1^+^ CD4^+^ T cells from either 3A9 or 5C.C7 TCR transgenic mice 3 days before the CFSE-labelled cohort. Draining lymph nodes from 4 mice per group were harvested 2.5 days after adoptive transfer of the CFSE-labelled cohort, and 3 mice from groups (i) and (iii) were sacrificed 8 weeks later for examination of draining lymph nodes and spleens. (b) Representative CFSE profiles of CD45.1^-^ 3A9 cells, gated for live CFSE^+^ CD4^+^ T cells. (c) Percentage of CFSE-labelled 3A9 TCR transgenic CD4^+^ T cells recruited into cell division. Each point represents an individual mouse and the group mean is indicated by the horizontal line. (d) Percentage of divided CFSE-labelled 3A9 TCR transgenic CD4^+^ T cells in each division peak. Each point represents the mean calculated from the 4 mice in each group and the error bars show the SEM. (e) Number of divided CD45.1^-^ CD4^+^ 3A9 TCR transgenic cells remaining in the draining lymph nodes and spleen in groups (i) and (iii) 8 weeks after immunisation. The number was calculated by multiplying the percentage of divided CD45.1^-^ CD4^+^ 3A9 TCR transgenic cells in each sample by the number of cells harvested from the organ. Each point represents an individual mouse and the group mean is indicated by the horizontal line.

### MHC Class II expression

Competition between MHC class I-restricted T cells has been linked to a T-dependent removal of specific peptide-MHC complexes from the surface of DCs, without an overall change in MHC expression per se (19). However in vitro studies have suggested an allele-specific downregulation of MHC class II after interaction between CD4^+^ T cells and APCs (20). Our specificity control was chosen to use a different MHC II allele so as to avoid antigen competition for presentation, and our model would therefore not have detected competition mediated by allele-specific downregulation. To test whether downregulation of MHC class II alleles occurs during the in vivo response, we measured the level of surface MHC Class II expression by DCs in the draining lymph nodes after immunisation with peptide-CFA. Intact 5C.C7 TCR tg mice in which 90% of CD4^+^ T cells were specific for MCC-IE^k^ were immunized to ensure that the majority of T cells and DCs within the node were involved in the response. Initially the level of both I-E^k^ and I-A^b^ expressed on the surface of DCs increased in mice immunised with MCC-CFA, compared with PBS-CFA (Fig. 8). MHC Class II expression declined subsequently and by day 7 was below the baseline level seen at day 0. However the expression of the allele presenting specific peptide to the majority of T cells (IE^k^) was higher in mice immunised with MCC-CFA, compared with PBS-CFA, whereas the level of IA^b^ was the same in the two groups. Thus we found no evidence of allele-specific downregulation, but rather saw a relative increase in the expression of the MHC allele presenting specific peptide to a large number of TCR transgenic T cells.

**Figure 8:**
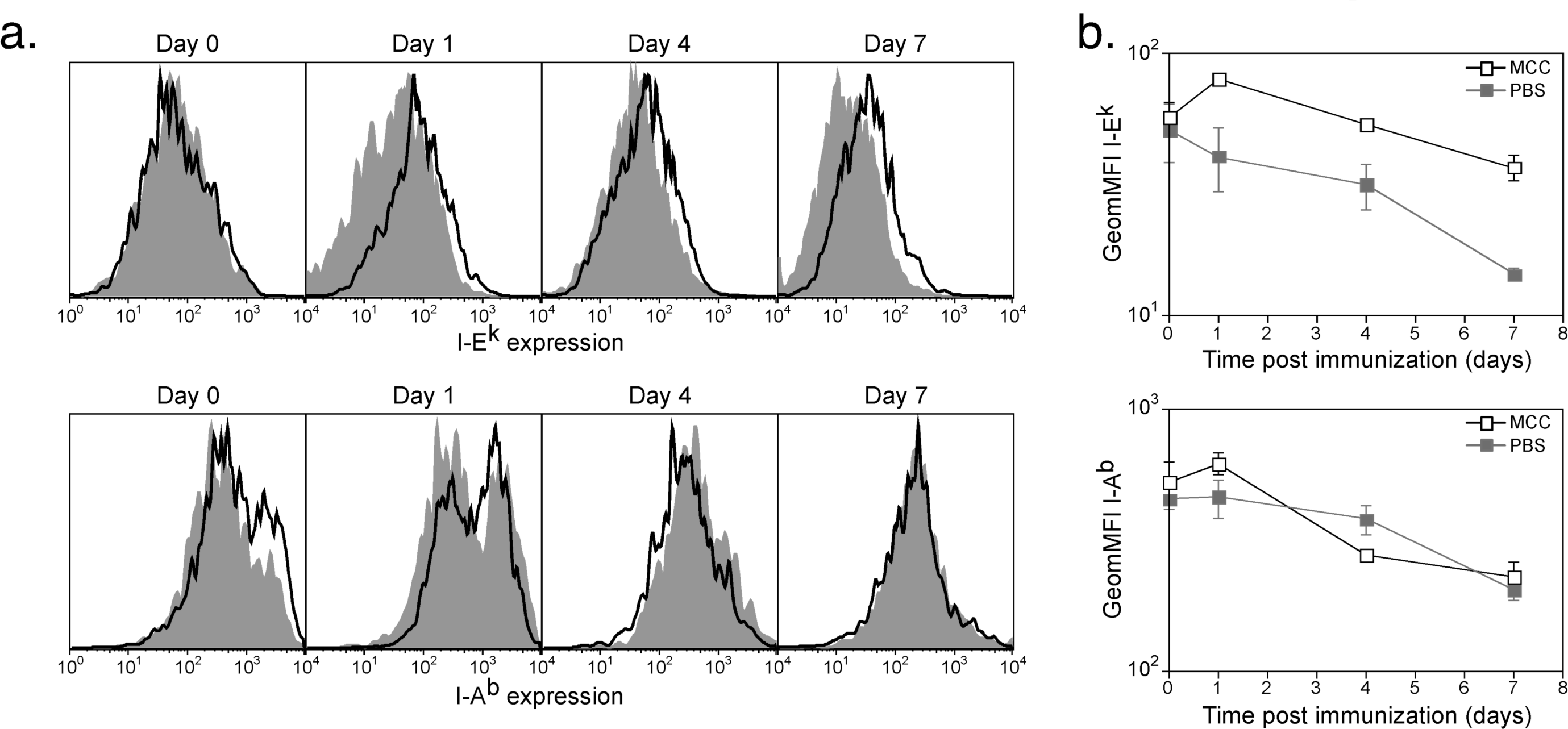
DC expression of MHC Class II molecules over the course of an adjuvant enhanced response. Intact 5C.C7 TCR transgenic mice were immunized with 10μg MCC peptide in CFA or PBS in CFA 1, 4 and 7 days prior to sacrifice. Draining lymph nodes harvested and the level of surface I-E^k^ and I-A^b^ was measured by flow cytometry. The equivalent lymph nodes were harvested from unimmunised mice to provide the day 0 data points. (a) Representative histograms showing surface expression of I-E^k^ (upper panel) and I-A^b^ (lower panel) by CD11c^+^, MHC Class II^+^ (M5-114^+^), B220^-^ DCs from mice immunized with MCC/CFA (unfilled profile) or PBS/CFA (filled profile). (b) Geometric mean fluorescence intensity +/- SEM of surface I-E^k^ (upper panel) and I-A^b^ (lower panel) expression on CD11c^+^, MHC Class II^+^ (M5-114^+^), B220^-^ DCs from 2 mice per group. Unfilled squares, MCC/CFA; filled squares, PBS/CFA.

## Discussion

We have shown here that recruitment of additional precursors into an ongoing response is an inverse function of the size of the response and is progressively inhibited with time (Figs 1-3). The process operates in a highly antigen specific manner to regulate the size of the response without affecting the response to unrelated antigens (Figs 4-7). Thus reduced recruitment into an ongoing response is not due to spatial constraints mediated by an increase in total number of activated cells, or an inability of naïve cells to access APCs. Recruitment is, however, a reflection of the level of antigen presentation and can be increased by administration of additional antigen at the time of T cell transfer (Fig. 3). In a response initiated by a small number of precursors, such as are present in the endogenous repertoire, a high proportion of exogenous naïve high affinity T cells can be recruited into the response up to 4 days after immunisation (Fig. 3). This is consistent with previous data in which a response of adoptively transferred TCR transgenic cells could be initiated up to three weeks after immunisation (21). However when the response is initiated with a relatively large number of high affinity precursors, recruitment of further cells into the response is significantly inhibited with 24 hours (Fig. 1). These data support the existence of a CD4^+^ T-cell dependent mechanism to control the ongoing level of specific antigen presentation, and thus to control the size of the primary immune response.

Interestingly both activated CD4^+^ (22) and CD8^+^ (23-25) T cells can continue to divide in vitro after removal of antigen. Memory cell differentiation is also believed to be independent of antigen-MHC (26, 27). However our results suggest that optimal proliferation and differentiation in vivo requires ongoing access to antigen-MHC (28). After acute activation leading to surface expression of CD69, not all activated cells are recruited into cell division within the next 2.5 days (as in groups (ii) and in the experiment shown in Fig. 4). Moreover, ongoing cell division and differentiation to cytokine secretion is even more profoundly affected by T cell competition than initial recruitment. For example, 40% of precursors were recruited in group (ii) in the experiment described in Fig. 2, compared with 70% in the control group (Fig. 2c). However the mean number of divisions per recruited cell was only 2.5, compared with >6 in the control (Fig. 2d). The number of IL-2 secreting cells was reduced from 10^6^ to 10^5^ (Fig. 2i) and IFN-γ-secreting cells dropped from 3×10^5^ to 2×10^4^ (Fig. 2n). While we were able to fully restore recruitment of additional precursors into division by the administration of additional antigen at the time of cell transfer (Fig. 3c, group (iv)), subsequent proliferation was still decreased compared to the control (Fig. 3d). Taken together, these results suggest that the level of available antigen not only controls initial T cell activation, but subsequent proliferation and differentiation of effector cells.

Previous studies in which antigen was administered by means of injecting peptide pulsed or virus infected DCs have suggested that antigen presentation in the context of MHC class I is relatively short-lived (29-31). The ability of CD8^+^ T cells to kill DCs presenting specific antigen is believed to contribute to the rapid loss of peptide-bearing DCs (32-34). In some studies, CD4^+^ T cells have also been shown to decrease the survival of DCs presenting specific antigen (35) whereas in others, CD4^+^ T cell recognition increased DC survival (36). Both CD4 and CD8 T cells have been shown to remove specific peptide-MHC complexes (37-39) from the surface of APCs. Indeed, Kedl et al. suggested that this process was responsible for antigen-specific competition between CD8^+^ T cells in vivo (19). Although we have no direct evidence that CD4^+^ T cells limit the size of the response by downregulating or removing cognate peptide-MHC complexes from the surface of DCs, it is likely that this mechanism is at least partly responsible for antigen specific competition between CD4^+^ T cells in vivo. Certainly we found no evidence for global downregulation of MHC Class II (Figure 8), consistent with the antigen-specific nature of T cell competition in this model. Future experiments will be required to measure the expression of specific peptide-MHC class II complexes on DCs during the course of the primary immune response.

Our data provides no support for an antigen non-specific competitive effect of *in vivo* immune responses, in contrast to previously published reports (17-19). Kedl et al. demonstrated a relatively small antigen non-specific effect, which could be overcome by injection of antigen pulsed DCs (17). They interpreted this result as supporting a model in which ongoing T cell responses deprive other T cells of access to DCs, globally suppressing other simultaneous responses. However, only a tiny fraction of injected DCs actually reach the T cell zone of the draining node, where they represent only 1-2% of total DCs (40). Moreover, we have shown that the injection of pulsed DCs can distribute Class II-MHC peptide complexes to other endogenous DCs, probably by means of exosomes (40) (Roediger, Chklovskaia and Fazekas de St. Groth, unpublished). Thus, the result of Kedl et al. could simply have reflected an increase in the concentration of specific peptide-MHC complexes, without any significant change in access to the DCs presenting them.

In our studies, a number of different manoeuvres were used to increase the sensitivity of the experimental model to possible competitive effects between cells directed to different MHC Class II restricted epitopes (Figs 5-7). Antigen doses were reduced so that cells would not be exposed to optimal concentrations, the readout was switched from the very high affinity 5C.C7 TCR to the lower affinity 3A9 TCR, the delay between the initiation of responses was increased to 4 days, so that the competing response was at its peak when the second response was initiated, and the size of the response was measured up to 8 weeks after priming. Under none of these conditions could antigen-non-specific competition be detected. To exclude the possibility that the two unlinked peptides were being presented by different DCs, a fusion protein requiring processing for presentation of both epitopes was administered, and gave the same results as the co-injection of a mixture of peptides.

In summary, the unique antigen specificity of competition between CD4^+^ T cells ensures that the immune system’s ability to respond to different antigenic epitopes is not compromised by an ongoing response to another antigen. The requirement for continued access to antigen for full proliferation and differentiation of CD4^+^ T cells ensures proliferation will continue in the presence of high levels of antigen but diminish once antigen levels decline. Together this provides a mechanism whereby the immune system responds efficiently to late phase antigens or secondary infections, while exerting tight control over the size and kinetics of each individual antigen specific response.

## Acknowledgments

The authors would like to thank Drs Didrik Paus and Robert Brink for adapting the HELMCC construct for expression in yeast, Dr Jennifer Kingham and the staff of the Centenary Institute Animal Facility for excellent animal husbandry, and Dr Adrian Smith and the staff of the Centenary Institute Flow Cytometry Facility for their expert assistance with flow cytometry.

## Grant Support

A.J.S. was supported by an Australian Post-Graduate Research Award. B.F.deSt.G. was supported by a National Health and Medical Research Council of Australia Principal Research Fellowship. This work was funded by a Program grant from the National Health and Medical Research Council of Australia. The support of the New South Wales Health Department through its research and infrastructure grants programme is gratefully acknowledged.

## Financial Disclosure

The authors have no conflicting financial interests.

## Abbreviations

CHO: chinese hamster ovarian,
DAPI: 2-(4-Aminophenyl)-6-indolecarbamidine dihydrochloride;
DC: dendritic cell;
HEL: hen egg lysozyme;
MCC: moth cytochrome C,
PI: propidium iodide.

## Notes

### Competing Interest Statement

The authors have declared no competing interest.

